# On the bioconversion of dietary carotenoids to astaxanthin and its distribution in the marine copepod, *Tigriopus californicus*

**DOI:** 10.1101/174300

**Authors:** Ryan J. Weaver, Paul A. Cobine, Geoffrey E. Hill

## Abstract

Red carotenoid-based coloration is widely distributed across marine and terrestrial animals and has taken a prominent role in studies of how phenotypic traits evolve in response to natural and sexual selection. Key to these studies is an understanding of the physiological mechanisms that give rise to red coloration, yet an ideal model system for such work has not been identified. The red marine copepod *Tigriopus californicus* is used as a model system for studies in ecotoxicology, genetics, and physiology, but the mechanisms involved in its bright red coloration have not been well studied. Like nearly all animals that display red carotenoid coloration, *T. californicus* likely convert yellow carotenoids present in their algal diet to red carotenoids. We conducted precursor/product feeding experiments to demonstrate that *T. californicus* bioconverts dietary carotenoids to the red carotenoid, astaxanthin. In separate treatment groups, copepods were fed carotenoids that are precursors to specific astaxanthin bioconversion pathways. We found that copepods from each precursor pigment group produced astaxanthin, and that the amount produced depended on which carotenoid was supplemented. We also describe the distribution of astaxanthin in developing egg sacs and show that the red color of the naupliar eyespot is not due to astaxanthin. We briefly discuss the potential of *Tigriopus californicus* to serve as a model system for the study of carotenoid metabolism in animals.

## 1. Introduction

With few exceptions, animals cannot synthesize carotenoids *de novo* from basic biological precursors (Britton and Goodwin 1982). To use carotenoids as external colorants, animals as diverse as flamingos and lobsters must obtain carotenoids from their diet (Fox and Hopkins 1966; Cianci et al. 2002). Animals with red coloration can derive red carotenoid pigments via two distinct pathways: they can ingest yellow pigments and oxidize them to produce red keto-carotenoids (Fig. 1) or they can ingest red pigments directly. For most metazoans, yellow carotenoids are much more common components of diets and thus most metazoans derive red coloration through the conversion of yellow dietary pigments (Goodwin 1984; Brush 1990). The processes involved in carotenoid bioconversion have far reaching implications for research across fields of study ranging from ecotoxicology (Weaver et al. 2016) to sexual selection for colorful ornaments (Hill 1991; Weaver et al. 2017). Yet, despite the long history of research on the evolution and distribution of carotenoid coloration across diverse taxa such as birds, fish, and crustaceans, an ideal model system for studying the genetic and physiological mechanisms involved in carotenoid metabolism in animals does not exist.

**Figure 1.**
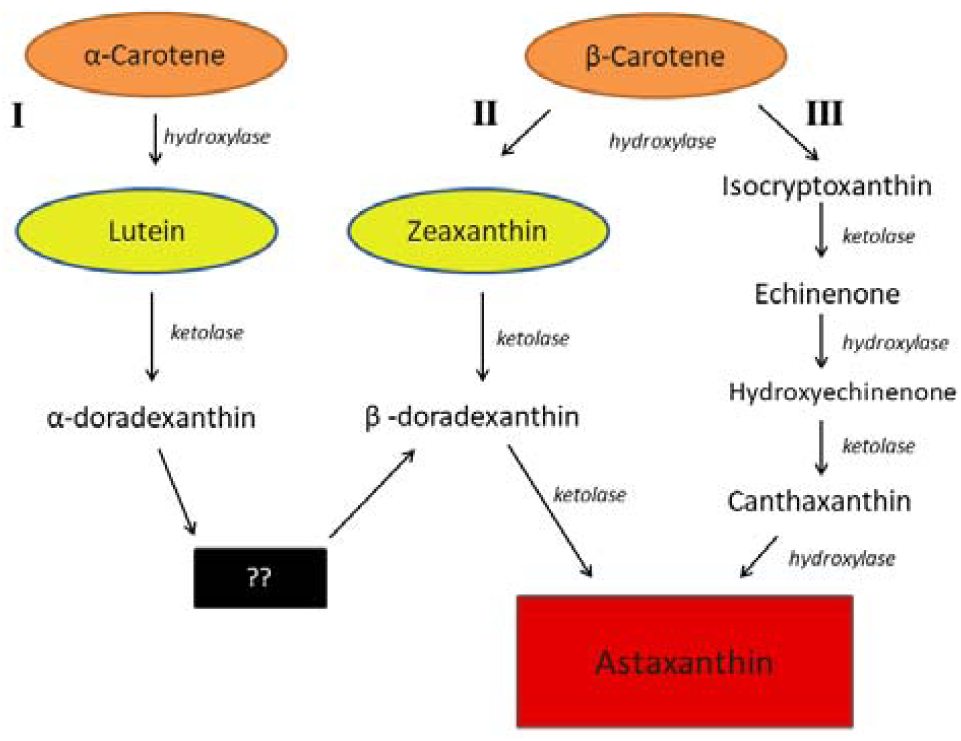
Proposed bioconversion pathways for astaxanthin production in animals (modified from Rhodes, 2007). Pathway I is used by some fishes, including goldfish, that utilizes lutein as a substrate. ?? represents the putative transformation of α to β-doradexanthin. Pathway II utilizes β-carotene or zeaxanthin as a substrate to astaxanthin. Pathway III begins with β-carotene and includes canthaxanthin as an intermediate to astaxanthin. The class of enzyme responsible for each transformation is italicized.

Recently, we have begun investigating the potential of the red marine copepod *Tigriopus californicus* (Baker 1912) to serve as a model system for the study of carotenoid physiology. Over the past four decades, *Tigriopus* copepods have become model organisms for studies on ecotoxicology (Raisuddin et al. 2007), phylogeography (Edmands 2001), local adaptation (Pereira et al. 2016), and mitochondrial-nuclear interactions (Ellison and Burton 2008). As a result, a wealth of physiological and genetic data exist that will facilitate detailed investigations in the molecular mechanisms involved in carotenoid coloration. However, fundamental aspects of their pigment physiology are relatively unexplored.

In the wild, *Tigriopus californicus* and other species within the genus *Tigriopus* typically have bright orange-red coloration (Fig. 2 a,c) that is produced via accumulation of the red keto-carotenoid astaxanthin (Goodwin and Srisukh 1949; Davenport et al. 2004; Weaver et al. 2016). The orange-red coloration is visible to the naked eye, but upon closer inspection under a dissecting microscope the pigment appears to be in the highest concentration along the gut lining and in lesser amounts throughout the exoskeleton. Adults retain a single naupliar eye spot located on the dorsal anterior cephalosome that is a vibrant red color (Fig 2). Previous studies have examined the microstructure of the naupliar eye and speculated that the red color was the result of carotenoids, but no formal analysis was undertaken (Martin et al. 2000). Carotenoids may also be deposited into eggs (Goodwin and Srisukh 1949). Wild gravid females and gravid females fed microalgae in the lab carry a single median egg sac of 15-40 developing embryos (Burton 1985) that transitions from a dark gray to rich red color as development progresses (Fig 2c).

**Figure 2.**
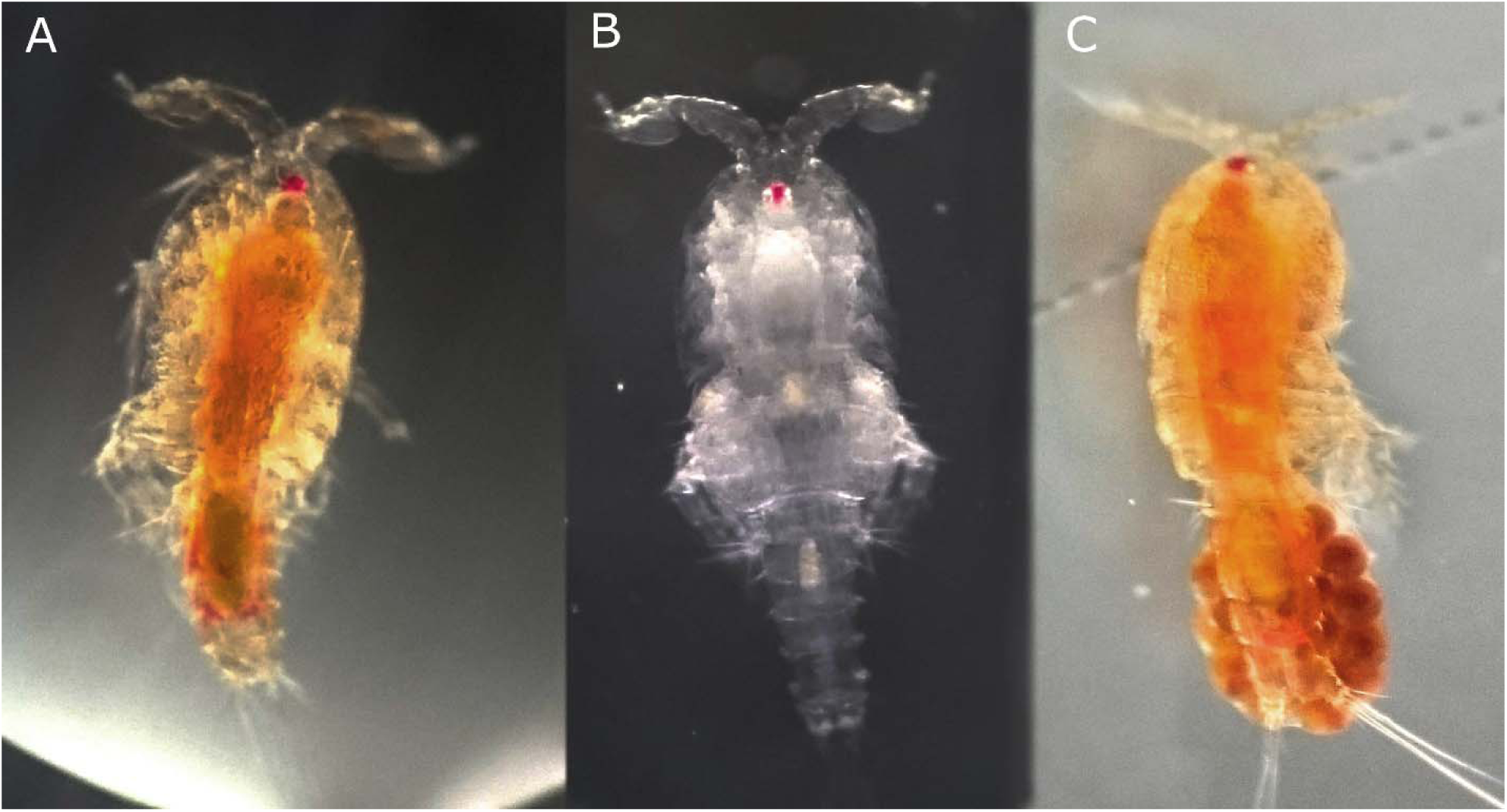
Coloration of *T. californicus* fed different diets. Typical coloration of *Tigriopus californicus* fed microalgae (A,C) and nutritional yeast (B). The red naupliar eye spot is present in copepods fed both carotenoid-rich and carotenoid-deficient diets suggesting that production of that eye pigment is not diet-dependent. (C) Females carry egg sacs that transition from dark gray to red as embryos develop.

The dietary source of carotenoid coloration of *Tigriopus* copepods is supported by the observation of Davenport et al. (1997) that when they are reared on a diet lacking dietary carotenoids, they lose their characteristic red color and appear white. However, to date, the dietary requirements and the bioconversion of dietary carotenoids to astaxanthin by *Tigriopus californicus* has not been the subject of well-controlled studies.

In this study, we first characterized the molecular source of pigmentation in *T. californicus*. We compared the body coloration and identified the carotenoid content of animals fed a live microalgae diet and carotenoid-free yeast diet. Next, we tested whether the red coloration of the naupliar eye spot (Fig. 2) and the coloration of the egg sacs of gravid females (Fig. 2c) was due to the presence of astaxanthin. Finally, we performed carefully controlled precursor-product carotenoid-supplement feeding experiments to study the bioconversion of dietary precursor carotenoids to astaxanthin by *T. californicus*.

## 2. Methods

### 2.1 Copepod culturing

*Tigriopus californicus* copepods collected from the wild in the vicinity of San Diego, California have been raised in our laboratory since 2014 in 10 L aquaria in filtered artificial seawater (ASW) at salinity of 32 at 24 °C on a 12 h light: 12 h dark light cycle and fed live microalgae *Tetraselmis chuii* and *Synechococcus spp.* This diet contains algal-derived carotenoids α- and β-carotene, lutein, and zeaxanthin (Guillard et al. 1985; Brown and Jeffrey 1992), which are hypothesized to be substrates for bioconversion to astaxanthin (Fig. 1). We refer to copepods raised under these conditions as our ‘stock population’.

### 2.2 Testing the dietary origin of red coloration in T. californicus

To test the hypothesis that the characteristic orange-red coloration of *T. californicus* depends on the presence of carotenoids in their diet, in 2015 we switched a sample of approximately 1,000 stock population copepods to a carotenoid-free diet of nutritional yeast (Bragg, Santa Barbara, CA). The nutritional yeast diet contains inactive dry yeast and a mix of B-complex vitamins but lacks carotenoids. We cultured these copepods in 15 L containers with opaque sides and lids to deter algal growth under gentle aeration; changing the water and mixing copepods from separate containers approximately once every three months. We refer to copepods raised under these conditions as ‘yeast-fed copepods’. We sampled 10 adult copepods from the stock population in triplicate and 10 yeast-fed copepods in triplicate then visually assessed coloration and tested for the presence of carotenoids in their bodies as described in section *2.6* below.

### 2.3 Qualitative carotenoid analysis of red eyespots

To determine whether the red coloration of the naupliar eyespot was carotenoid-based, we dissected the portion of the cephalosome containing the eyespot from 10 adult yeast-fed copepods. All 10 dissected eyespots were pooled in one tube and the corresponding 10 bodies were pooled in a separate tube then processed and analyzed for carotenoids as described below.

### 2.4 Maternal deposition of astaxanthin to developing embryos

In a separate experiment, we sought to determine whether the red coloration of mature egg sacs attached to stock population gravid females was carotenoid-based. We carefully removed red egg sacs from three gravid females using a fine needle under a dissecting scope and processed each clutch individually for carotenoid analysis as described below.

### 2.5 Bioconversion to astaxanthin from carotenoid supplementation of yeast-fed copepods

To definitively test for the bioconversion of dietary carotenoids to astaxanthin by *T. californicus,* yeast-fed copepods were supplemented with either β-carotene, lutein, zeaxanthin, and canthaxanthin. We chose these four precursor carotenoids for our feeding experiments because they occupy various positions in the putative pathway used by copepods to produce astaxanthin (Fig. 1). Stock solutions of β-carotene, lutein, zeaxanthin, and canthaxanthin were made from water-soluble carotenoid beadlettes (DSM, Basel, Switzerland) in ASW and diluted to a working carotenoid concentration of 2 μg mL^−1^. Each carotenoid-supplement contained only the carotenoid listed, except the lutein supplement. Of the total carotenoid content, 90.6% was pure lutein, but zeaxanthin comprised 7.8%. Therefore, the “lutein” supplement contained 1.984 μg mL^−1^ lutein, and 0.016 μg mL^−1^ zeaxanthin. For each carotenoid supplement group, 10 adult copepods were placed in 5 mL carotenoid solution in each well of a six-well plate (*n* = 6 for each supplement group) with 0.75 mg nutritional yeast as food for 48 h, then processed for carotenoid analysis (see below).

### 2.6 Carotenoid analysis

After each experiment, copepods were placed in fresh ASW to clear gut contents, then rinsed with de-ionized water, dried and –for the bioconversion experiment – weighed to the nearest 0.01 mg. The mass from one sample from the β-carotene supplement group was not recorded and one sample from the zeaxanthin supplement group was destroyed before carotenoid analysis.

Carotenoids were extracted from copepods by sonicating in 500 μL HPLC-grade acetone in a 1.7 mL microcentrifuge tube for 10s at 10W; we then capped tubes with nitrogen gas and incubated them overnight at 4 °C in the dark. Samples were centrifuged at 3,000g for 5 min, the supernatant was removed to a new tube and evaporated to dryness at 40 °C under vacuum, then resuspended in 50 μL acetone. Carotenoids were separated using a Shimadzu HPLC system from a 40 μL injection on to a Sonoma C18 column (10 μm, 250 x 4.6 mm, ES Technologies) fitted with a C18 guard cartridge. We used mobile phases A) 80:20, methanol: 0.5 M ammonium acetate, B) 90:10, acetonitrile:H_2_O, and C) ethyl acetate in a tertiary linear gradient as follows: 100% A to 100% B over 4 min, then to 80% C: 20% B over 14 min, back to 100% B over 3 min and returning to 100% A over 5 min and held for 6 minutes (Wright et al. 1991). Total run time was 32 min at a flow rate of 1 mL min ^−1^. Absorbance was measured at 450 nm using a UV/VIS detector. Carotenoids were identified and quantified by comparison to authentic standards. Astaxanthin concentration was normalized to copepod dry weight.

### 2.7 Statistical analysis

We tested for a difference in the amount of astaxanthin produced by copepods from each group using ANOVA and evaluated pairwise comparisons between groups using Tukey HSD post-hoc test.

## 3. Results

### 3.1 Dietary origin of the red coloration of T. californicus

The major carotenoid found in stock population copepods was free astaxanthin (mean ± SE; 49.38 ± 2.19 ng copepod ^−1^, *n =* 3). Minor amounts of mono and di-esterified astaxanthin were detected comprising 3.22% and 8.83% of the total carotenoid content, respectively. We found that when stock population copepods were switched to a carotenoid-free yeast diet they lost their characteristic orange-red coloration and appeared clear (Fig. 2b). Biochemical analysis revealed that a small but measurable amount of astaxanthin was detected in the yeast-fed copepods (mean ± SE; 0.54 ± 0.016 ng copepod ^−1^, *n* = 3).

### 3.2 Astaxanthin analysis of red eyespots

We detected no carotenoids in the red eyespot-enriched fraction of yeast-fed *Tigriopus californicus.* Analysis of the corresponding body fraction returned a similar concentration of astaxanthin as from the yeast-fed copepods in the previous experiment (0.3 ng copepod ^−1^).

### 3.3 Astaxanthin analysis of red egg sacs

We found that the red egg sacs of stock population gravid females contained astaxanthin (mean ± SE; 10.53 ± 3.31 ng egg sac ^−1^, *n* = 3).

### 3.4 Bioconversion of dietary supplemented carotenoids

We found that *T. californicus* copepods from each carotenoid supplement group accumulated astaxanthin after 48 h when none was present in their diet (Fig. 3). The amount of astaxanthin converted depended on the specific carotenoid supplemented (mean astaxanthin in μg mg ^−1^ dry mass of copepod ± SE); zeaxanthin (0.99 ± 0.11 μg mg ^−1^, *n* =5) > canthaxanthin (0.90 ± 0.12 μg mg ^−1^, *n* =6) > β-carotene (0.38 ± 0.06 μg mg ^−1^, *n* =5) > lutein (0.21 ± 0.02 μg mg ^−1^, *n* =6). We detected significant differences in astaxanthin production among supplement groups (ANOVA, *F* (4,23) = 29.83, *P* < 0.0001). Post-hoc pairwise comparisons revealed that the amount of astaxanthin produced was not significantly different between zeaxanthin and canthaxanthin groups (Tukey HSD, difference, (95% CI): −0.09, (−0.41 to 0.23) μg mg ^−1^, *P=*0.91) or between β-carotene and lutein groups (Tukey HSD, difference, (95% CI) : −0.17, (−0.5 to 0.14) μg mg ^−1^, *P=*0.49). However, we found that copepods supplemented with zeaxanthin and canthaxanthin produced significantly more astaxanthin than copepods fed either β-carotene or lutein (*P < 0.001*). In each supplement group, only free astaxanthin and the supplemented carotenoid were detected. It is possible that exiguous amounts of intermediate carotenoids were present in samples, but were not detected on our system.

**Figure 3.**
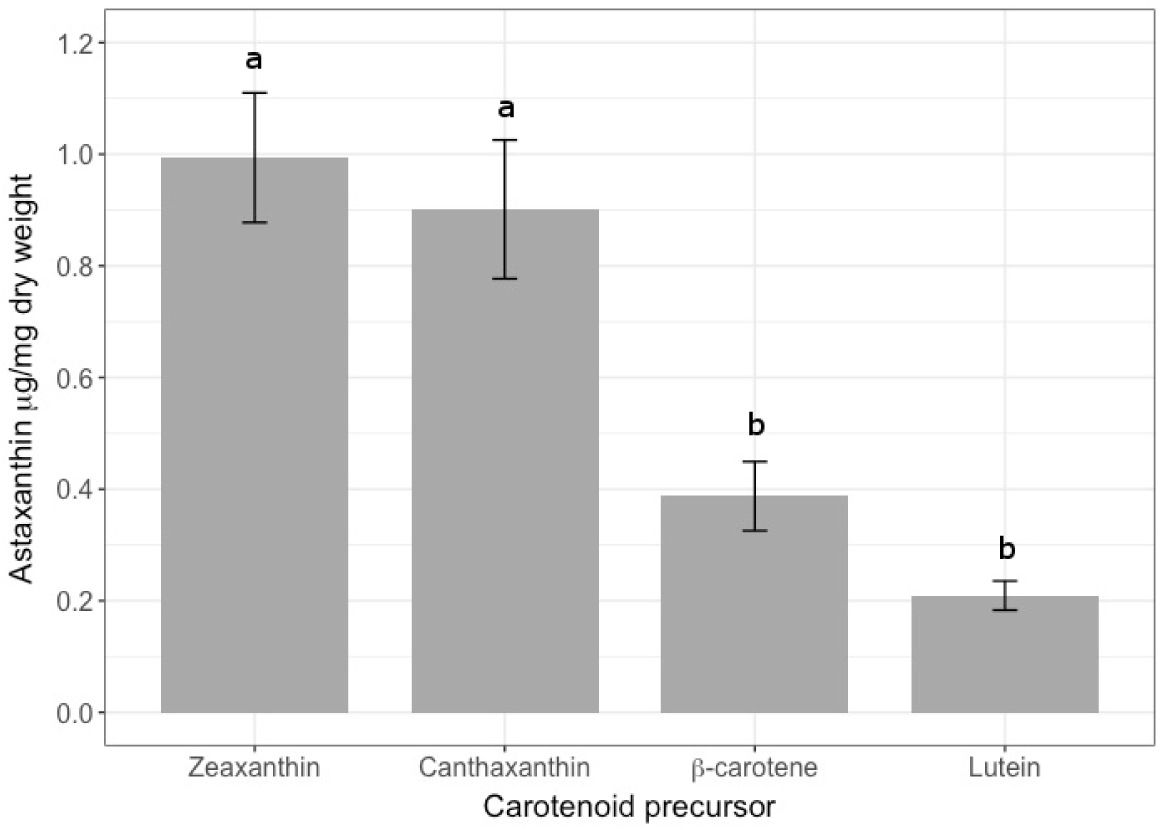
Bioconversion of dietary carotenoids. Astaxanthin content of *T. californicus* copepods after supplementation with a precursor carotenoid upstream of astaxanthin for 48h. Least-squared means and standard error bars are shown and significantly different means are indicated by separate letters.

### 4. Discussion

In this study, we conducted carefully controlled precursor/product tests to document the bioconversion of dietary carotenoids to the red ketocarotenoid astaxanthin in the marine copepod *T. californicus*. When they were maintained on a yeast diet that provided no carotenoids, copepods became clear with no hint of red or yellow. Biochemical analysis confirmed that these animals essentially lacked carotenoids in their tissues. When we supplemented yeast-fed carotenoid-free copepods with β-carotene, lutein, zeaxanthin, or canthaxanthin for 48 h, we observed significant production of astaxanthin (Fig. 3). These experiments confirmed that *T. californicus* requires carotenoid-rich foods to obtain their characteristic orange-red color and that they bioconvert precursor carotenoids from their diet to astaxanthin.

The amount of astaxanthin produced depended on which carotenoid was the precursor for bioconversion (Fig 3). Copepods fed zeaxanthin and canthaxanthin produced more astaxanthin in 48h than copepods fed β-carotene or lutein. This pattern suggests that the number of oxidation reactions required to convert the supplemented precursor carotenoid to astaxanthin may mediate the rate of astaxanthin production in this species. Canthaxanthin and zeaxanthin require two hydroxylation or two ketolation reactions respectively to form astaxanthin; whereas, β-carotene requires 4-7 reactions, and lutein requires 3 reactions and a conversion from α-doradexanthin to β-doradexanthin (Fig 1). Interestingly, a similar effect of supplementation with zeaxanthin versus lutein was observed in a study of American Goldfinches, which transform dietary carotenoid to Canary Xanthophyll A and B, and Northern Cardinals which transform dietary pigments to astaxanthin. Both goldfinches and cardinals produced more oxidized pigments and more colorful integumentary structures when they were fed zeaxanthin compared to when they were fed lutein (McGraw et al. 2014).

In the wild, copepods that feed on micro and macro algae ingest relatively large quantities of β-carotene, zeaxanthin, and lutein (Brown and Jeffrey 1992; Buffan-dubau et al. 1996; Sigaud-Kutner et al. 2005; Takaichi 2011; Wang et al. 2015). The bioconversion of dietary carotenoids to astaxanthin has been documented in other copepod species (Rhodes 2007; Caramujo et al. 2012), crustaceans (Hsu et al. 1970; Tanaka et al. 1976) and fish (Hsu et al. 1972) and these authors have concluded that the pathway begins with β-carotene. However, in addition to being used as a pigment, β-carotene is also the main precursor for vitamin A synthesis in animals (Parker 1996), which may cause an allocation tradeoff between vitamin A production and coloration (Hill and Johnson 2012). Alternatively, our results suggest that *T. californicus* may utilize multiple carotenoids as substrates for bioconversion to astaxanthin depending on which carotenoids are available in their diet, and/or the body’s need for vitamin A. Zeaxanthin has no vitamin A capacity and we found that copepods fed this precursor produced significantly more astaxanthin than β-carotene supplemented copepods. While we did not detect any intermediates along the proposed bioconversion pathways, our results demonstrate that *T. californicus* use zeaxanthin as a substrate for astaxanthin production. It is possible that zeaxanthin is the start of a more efficient bioconversion pathway for astaxanthin production by *T. californicus.* Future experiments that analyze larger amounts of copepods within shorter sampling intervals may identify intermediate carotenoids and help resolve which astaxanthin bioconversion pathway(s) is used by *T. californicus*.

It is unclear from this study whether *T. californicus* uses lutein as a substrate for astaxanthin production because the lutein supplement also contained trace amounts of zeaxanthin. Lutein is common in marine phytoplankton as well, and some marine animals are thought to preferentially use lutein as the substrate for bioconversion to astaxanthin and other ketocarotenoids. Hsu (Hsu et al. 1972) and Katayama (Katayama et al. 1973) have shown that goldfish (*Carassius auratus*) potentially use lutein as precursor to astaxanthin. However, this bioconversion pathway requires the isomerization of α-doradexanthin to β-doradexanthin, a transformation that others have shown to be unlikely in fungus, plants, and other marine animals (Matsuno et al. 1999; Ohkubo et al. 1999).

The red coloration of the eyespot is not dependent on diet; *Tigriopus* copepods fed either algae or yeast both have red eye coloration (Fig 2). We found that the bright red eyespot coloration of *T. californicus* is not from astaxanthin, or any other carotenoid that we are able to detect, and that the trace astaxanthin content of yeast-fed copepods is not located in the eye. These results clarify that the red eye coloration is not from astaxanthin, but may be from the visual pigment rhodopsin that may utilize 3-hydroxyretinal as the chromophore (Cronin 1986).

We found that females deposit astaxanthin to developing embryos, supporting previous reports of this ketocarotenoid occurring in the egg sacs of other *Tigriopus* species (Goodwin and Srisukh 1949). It has been proposed that deposition of astaxanthin to developing eggs provides embryos photoprotection from solar UV radiation (Dethier 1980). Won et al. (2014) have shown that experimental UV exposure reduced hatching success of *Paracyclopina nana* nauplii, although the specific roles that carotenoids may play in survival following UV exposure remain unclear.

Investigations into the genetic architecture and physiological mechanisms involved in carotenoid metabolism in animals has only recently begun. The gene responsible for ketolation of yellow dietary carotenoids in birds - dubbed the redness gene - was independently discovered by Lopes et al (2016) and Mundy et al (2016). This gene encodes a cytochrome P450 oxidoreductase enzyme, CYP2J19, that has sequence motifs that implicate subcellular localization to mitochondria. The enzyme that enables conversion of yellow to red pigments in arthropods in general and *Tigriopus* copepods in particular has yet to determined, but *Tigriopus* are poised to be the model for identification of the physiological mechanisms and cellular locations involved in hydroxylation and ketolation of carotenoids in animals.

### 5. Conclusion

We have demonstrated that the orange-red coloration of *Tigriopus californicus* is a result of bioconverting dietary yellow carotenoids to the red keto carotenoid astaxanthin. Our experimental design of producing carotenoid-deficient copepods then supplementing with a single precursor carotenoid could serve as a powerful system for elucidating the genetic underpinnings and physiological constraints of carotenoid metabolism in this species, with broad implications applicable to understanding the evolution of carotenoid-based ornaments in honest signaling systems.

### 6. Compliance with Ethical Standards

The authors declare that they have no conflict of interest and that this research was conducted in compliance with all United States and Auburn University regulations on the use of invertebrate animals in research.

## 7. Acknowledgements

The authors thank Ron Burton and Felipe Barreto for advice on culturing conditions and the Hill and Hood lab groups for feedback on previous drafts of this manuscript.

